# Towards a bionic IoT: environmental monitoring using smartphone interrogated plant sensors

**DOI:** 10.1101/2022.03.10.483755

**Authors:** Yunlong Guo, John Canning, Zenon Chaczko

## Abstract

The utilisation of plants directly as quantifiable natural sensors is proposed. A case study measuring surface wettability of Aucuba japonica, or Japanese Laurel, plants using a novel smartphone field interrogator is demonstrated. This plant has been naturalised globally from Asia. Top-down contact angle measurements map wettability on-site and characterise a range of properties impacting plant health, such as aging, solar and UV exposure, and pollution. Leaves at an early age or in the shadow of trees are found to be hydrophobic with contact angle *θ* ~ 99°, while more mature leaves under sunlight are hydrophilic with *θ* ~ 79°. Direct UVA irradiation at *λ* = 365 nm is shown to accelerate aging, changing contact angle of one leaf from slightly hydrophobic at *θ* ~ 91° to be hydrophilic with *θ* ~ 87 ° after 30 min. Leaves growing beside a road with heavy traffic are observed to be substantially hydrophilic, as low as *θ* ~ 47°, arising from increased wettability with particulate accumulation on the leaf surface. Away from the road, the contact angle increases as high as *θ* ~ 96°. The results demonstrate that contact angle measurements using a portable diagnostic IoT edge device can be taken into the field for environmental detection, pollution assessment and more. Using an internet connected smartphone combined with a plant sensor allows multiple measurements at multiple locations together in real-time, potentially enabling tracking of parameter change anywhere where plants are present or introduced. This hybrid integration of widely distributed living organic systems with the internet marks the beginning of a new bionic internet-of-things (*b*-IoT).

## Introduction

Sensors are transforming the internet from a passive to an active smart internet-of-things (IoT) network system, or systems, with future global reach that have the capacity to transform society. For example, globally tracking pandemics in real-time [1] can transform health to quickly suppress outbreaks. As the full potential grows our understanding of the IoT, including the concept of smart cities, needs revaluation from the way consumers engage to how society functions [2]. The management of such systems involves long term developments in signal processing, both passive and active, potentially opening genuine artificial intelligence in time. However, traditional sensors in IoT are not always robust nor reliable long term, and their fabrication involves substantial deleterious climate impact that has largely been ignored. An exception has been the significant effort to develop self-powered devices that can operate indefinitely. Despite these efforts, the sensor itself remains unchanged in concept - a low-cost, traditionally manufactured simple structure often targeting an individual parameter. In this paper, we offer a different approach to sensing that foresees a future of climate friendly environmental monitoring using instead natural sensors that can, in principle, be integrated into the network. These bring additional benefits, including health and wellbeing for the wider community and environment [3]. To demonstrate this, a common plant species across Asia, aucuba japonica (or Japanese Laurel), which has also become naturalised in many locations elsewhere, is explored as an environmental sensor. Aucuba japonica, a popular ornamental shrub in China, is a compact evergreen plant with green-yellow variegated leaves [4] used in natural medicine [5]. It was introduced in the southern UK from Japan in 1783 and found to have recognised in woodlands by 1981 [6,7]. The high global prevalence of this plant, including in Australian subtropical zones in the southern hemisphere [8–12], makes it a potential candidate for a distributed sensing network that can help address global challenges such as climate change.

Plants make ideal practical, low-cost sensor alternatives to contemporary physical hardware traditionally used to monitor a range of environmental parameters such as air quality and health. Of interest is the robustness of this plant over time. Aucuba japonica has a relatively thick top cuticle layer, composed of cutin, a wax-like hydroxy fatty acid that protects the plant from direct UV irradiation and retains water internally whilst keeping the surface relatively dry by repelling it. In humid climates, these properties play an important role in plant robustness and keeping it free from disease. That, in turn, makes them the ideal choice for sensor evaluation. Aucuba japonica’s value as a specific living sensor coupled with a smartphone diagnostic is proposed and used to monitor the immediate surrounding environment. Such integration of living and non-living technologies can form the beginnings of what can be described as a bionic internet-of-things, or *b*-IoT.

With the selected sensor, the key technology component becomes the choice of portable and remote operating diagnostic. To recognise further the potential of this approach, this work focuses on using a widely available consumer product, the smartphone [15,16], which can allow general community access to diagnostics. This simply recognises that practical environmental monitoring requires as much social and community engagement as technology and insight, and so a future is envisaged that develops accessible natural and “green” technology to foster such engagement. The underlying principles demonstrated here have significant ramifications for policy more broadly in other sectors, including industrial and agriculture – simply having healthy plants on site can greatly relax monitoring demands for all site workers, for example. On agricultural land, plants offer other enormous benefits in recycling soil virility over time and improving local climate. The possibility exists of using crops themselves in a similar fashion for sensing, a novel form of agricultural or farm-based sensing. Importantly, plants generally bring aesthetic and wellness features both for humans and the environment more broadly.

In a well-known example, plant colour changes when pH changes, especially in the presence of Al which raises soil toxicity [17]. There are many other plant-derived parameters that can provide ongoing information of more subtle changes in a sensor format. Monitoring Aucuba japonica’s leaf surface properties can provide information about the plant’s health, its rate of photosynthesis and as well the surrounding environment whilst being well away from toxicity [18,19]. In particular, there are reports that leaf surface wettability is affected by air pollutants [20,21]. By extension, this would suggest that knowing contact angle (CA or *θ*) values can provide information and insight on the surrounding environment over time. However, the use of CA for this purpose is not practically feasible because conventional side view CA measurements are restricted to benchtop instruments requiring samples to be brought to a laboratory. Real or near real-time measurements are currently not feasible. Importantly, direct side view measurements are limited to sample edges and often will not correlate with the interior surface of a leaf [22]. Transport and cutting of the leaf imply a need for the preservation of conditions, limiting broader mapping of wettability. This has confined the study of plants and other organisms to curious scientific interests. In this work, this approach is disrupted, showing how surface wettability of distributed plants, if applicable in the field, can be a powerful tool for environmental monitoring. Coupled with smart devices such as smartphones, a powerful new bionic sensing approach can address shortcomings, including sensor limits, reliability, and sustainability in current IoT technologies for environmental monitoring.

To demonstrate proof of concept, a simple smartphone-based top-down CA measurement instrument was developed, taking the previous laboratory-based topdown contact angle mapping [23] into the field. Measuring and mapping the CA across a whole surface can be carried out easily by imaging multiple droplets either in quick succession or simultaneously. This addresses all issues with current instruments. Its potential is demonstrated by studying Aucuba japonica, comparing young and old leaves under various conditions, studying the exposure of the leaf to UV light, and assessing the effect of local environmental pollution from vehicles.

## Experimental section

### Instrument and measurements

Measuring the wettability of organisms can be challenging. For leaves this is because there is surface irregularity arising from varying curvatures, roughness, absorption, and reflection from its unique cuticle structure, features that make side measurements of contact angle unreliable. To access the interior of the leaf involves cutting samples into pieces. Top-down CA imaging (where the drop is imaged from the top and the CA extracted from diameter measurements) can circumvent these problems and map CA distribution over the actual leaf, providing substantially more information and data about a leaf’s overall wettability. To enable this in the field, a smartphone is used as the image capturing and measurement tool for on-the-spot diagnostics, taking CA measurements out of the laboratory. Importantly, using a connected device such as a smartphone opens opportunities to later couple multiple devices in various locations – so-called swarm technologies such as those proposed for addressing pandemics [1]. Cloud analytics can simultaneously collect, integrate, and analyse the data to provide additional information and insight that can be sent back to multiple users. The advantage of networked sensors is large data generation that can help circumvent challenges involved with less accurate individual measurements – these networked super instruments generate resilience to individual instrument error. This will form the basis of novel, hybrid organic-inorganic bionic IoT.

### The Contact Angle Analyser

The CA analyser prototype constructed in this work is illustrated in Fig 3. A smartphone (Model: Meizu 17) with a camera (SONY IMX686 COMS, 64 MP) is used to capture both top-down and side-view images of droplets so that they can be directly compared. A small aperture with a certain diameter was placed beside the sample for imaging calibration within the algorithm used to calculate the CA. A homemade attachment on which the smartphone sits (dimensions *x,y,z* = 114 × 97 × 128 mm) together with a protruding structure (dimensions *x,y,z* = 110× 109 × 98 mm), is designed to allow both side and top-down imaging of a drop at the same location.

**Fig 1.**
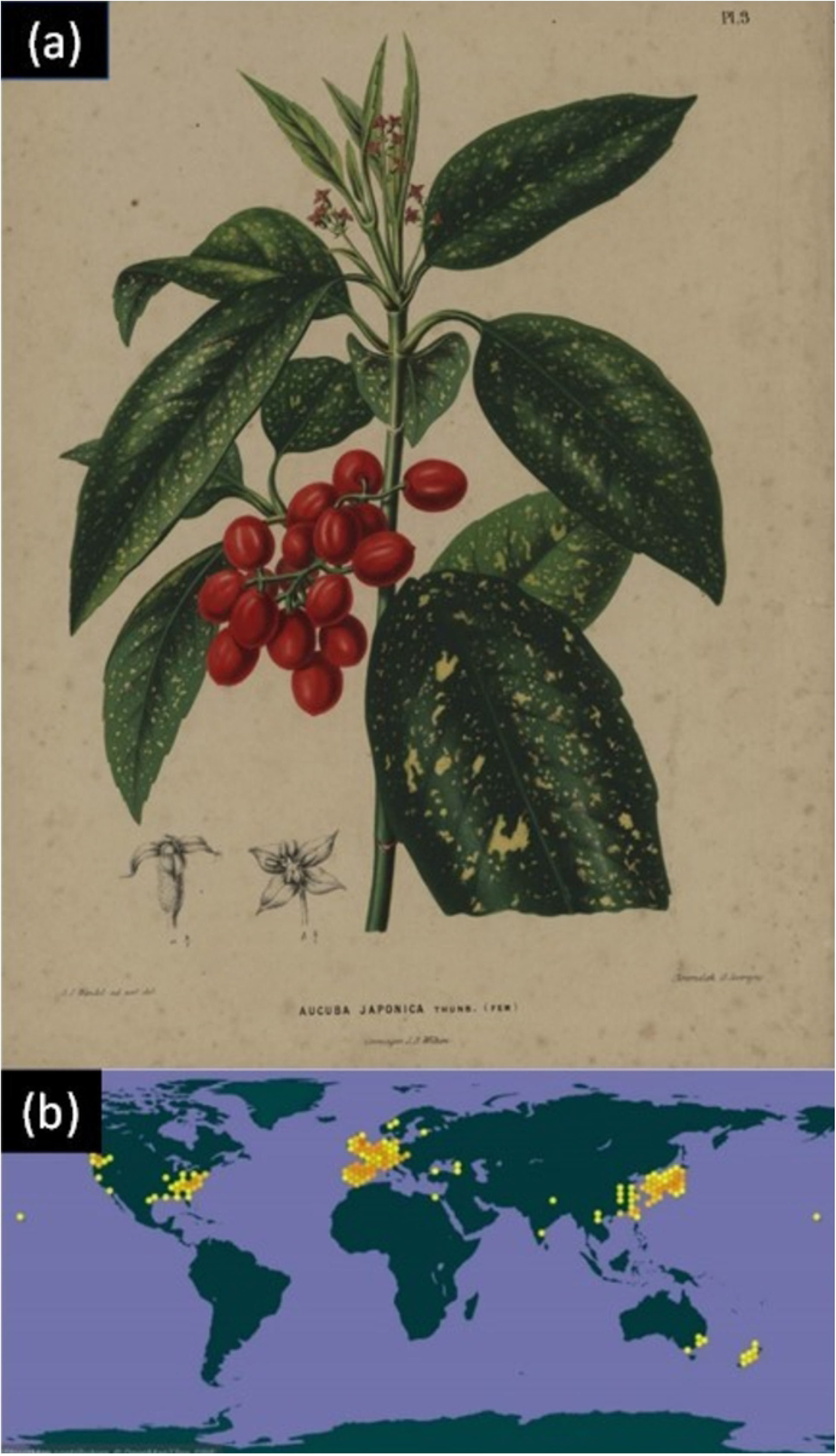
(a) Painting by Dutch artist A.J. Wendel, 1868. Reproduced from [13]; (b) Distribution of Aucuba japonica throughout the world mapping the potential coverage of a natural sensor network. The data was updated on 29/12/2021 by GBIF.ORG [14].

**Fig 2.**
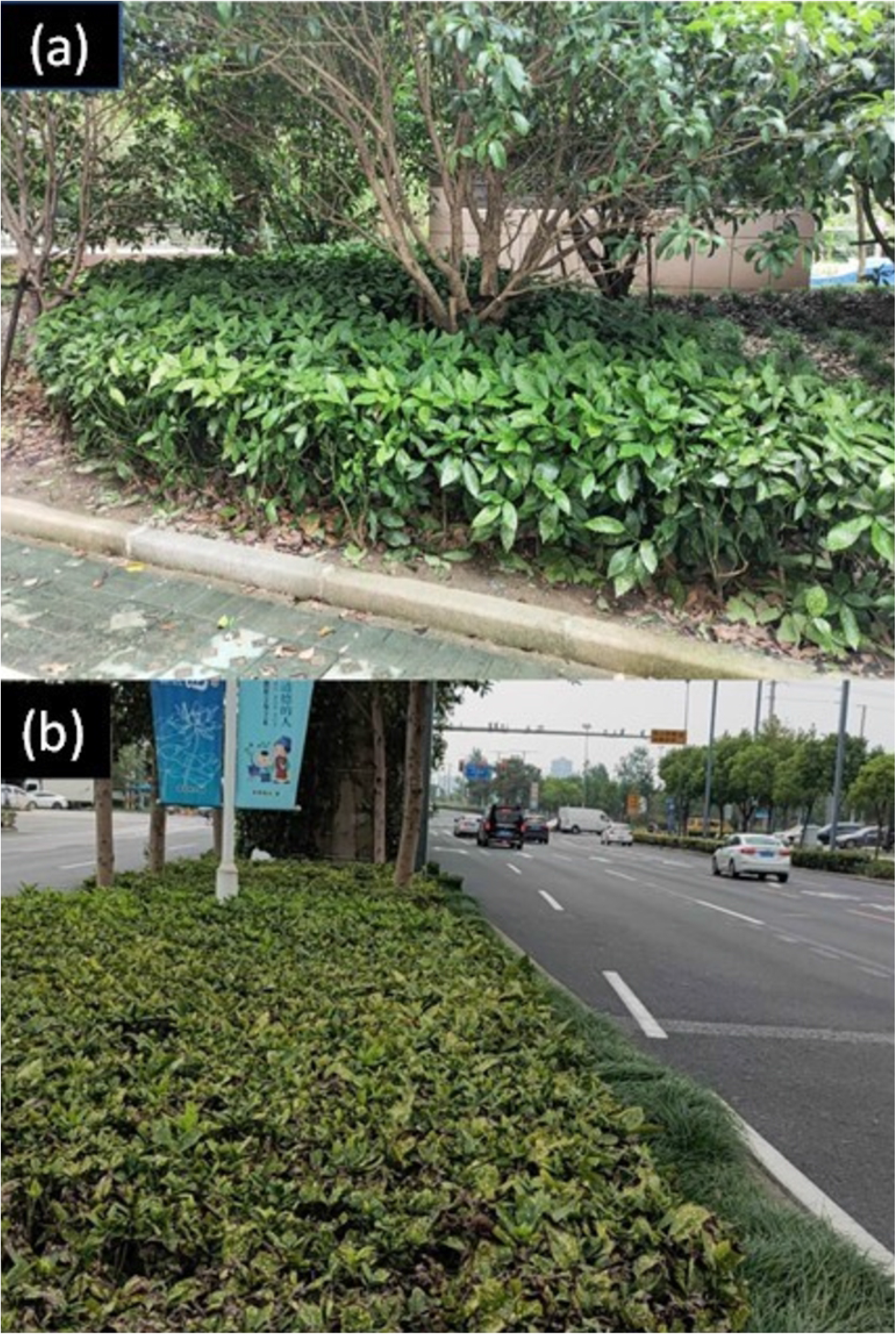
Image of Aucuba japonica: (a) plants sheltering under a tree away from a busy street and (b): plants next to a busy street. A striking difference is observed in yellow coloration and general plant vigor related to their local environment. The plants are exposed to varying conditions, including particulate matter released from vehicle exhaust, sun, wind, and rain.

**Fig 3.**
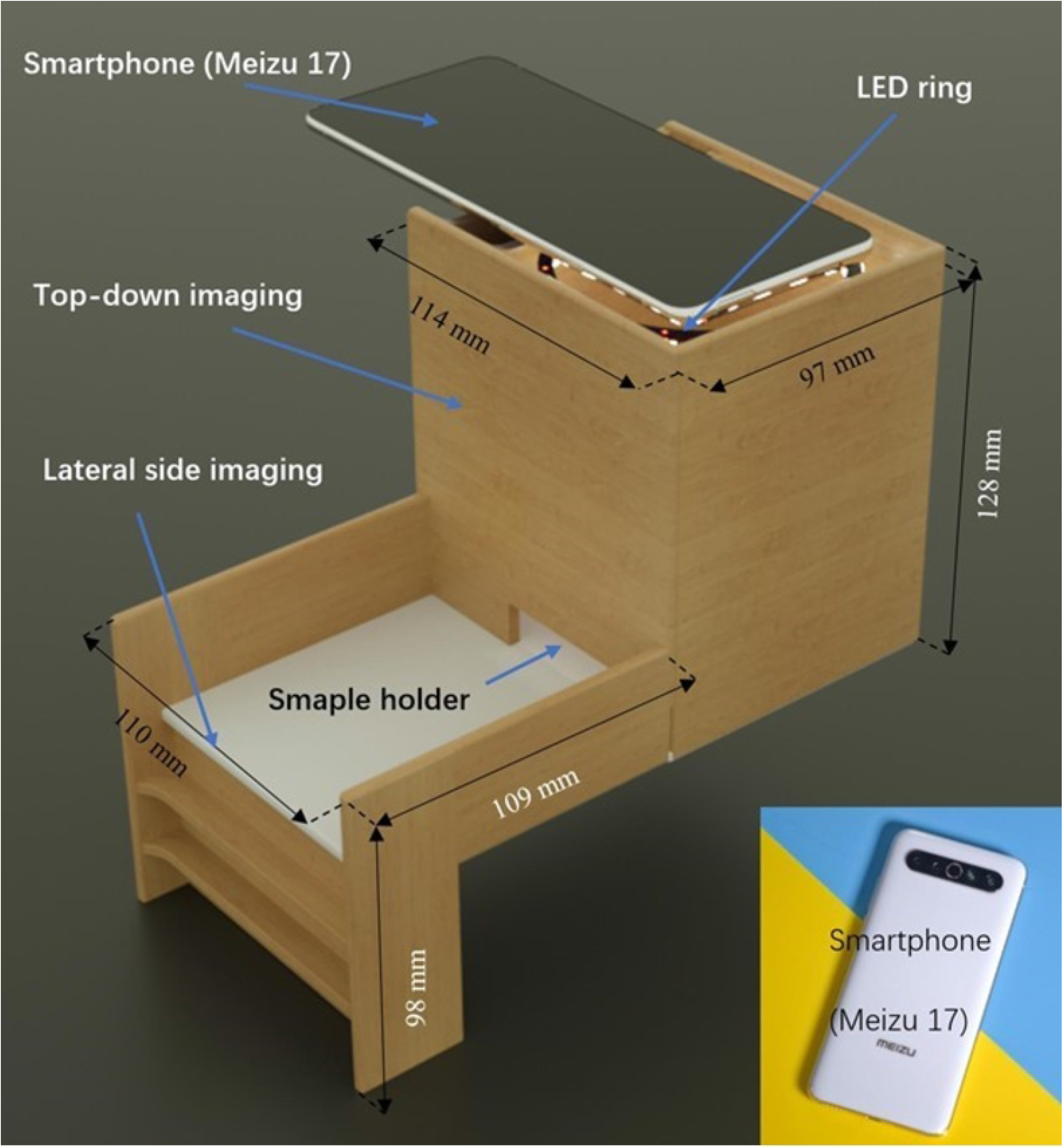
3D model of custom attachment. Top-down imaging occurs at the top using a white light-emitting diode (LED) ring to uniformly illuminate the drop evenly from the top. Inset shows an image of the smartphone used in this work.

A pipette (*V* = 0.5 to 10 *μ*L; accuracy ± 5%) is used to create the droplets that have a volume *V* = (4 ± 0.2) *μ*L. Drops at this small volume ensure that surface tension is greater than the gravitational force so that an approximate spherical-shaped caplet is obtained. From the diameter of this caplet, the contact angle can be readily derived in both hydrophilic and hydrophobic regimes (or omniphilic and omniphobic for liquids other than water).

A ring-shaped LED white light source is used to uniformly illuminate the samples. The smartphone LED, and torch functions are intentionally avoided because they are offset from the smartphone CMOS detector, which will generate drop shadows at the bottom of the attachment, effectively decreasing the CA values measured without additional algorithmic post-processing. A ring-shaped LED light offers superior illumination compared to a single LED light that requires the drops to be precisely centred to avoid shadowing. The configuration used is found to give robust and reliable measurements.

### Contact angle calculation

The diameter, *ϕ* =2*r*, of the spherical caplet segment in a top-down image, can reflect one of two different situations, shown in Fig 4. Hydrophilic surfaces have a CA lower than *θ* = 90° and hydrophobic surfaces have a CA higher than *θ* = 90°.

**Fig 4.**
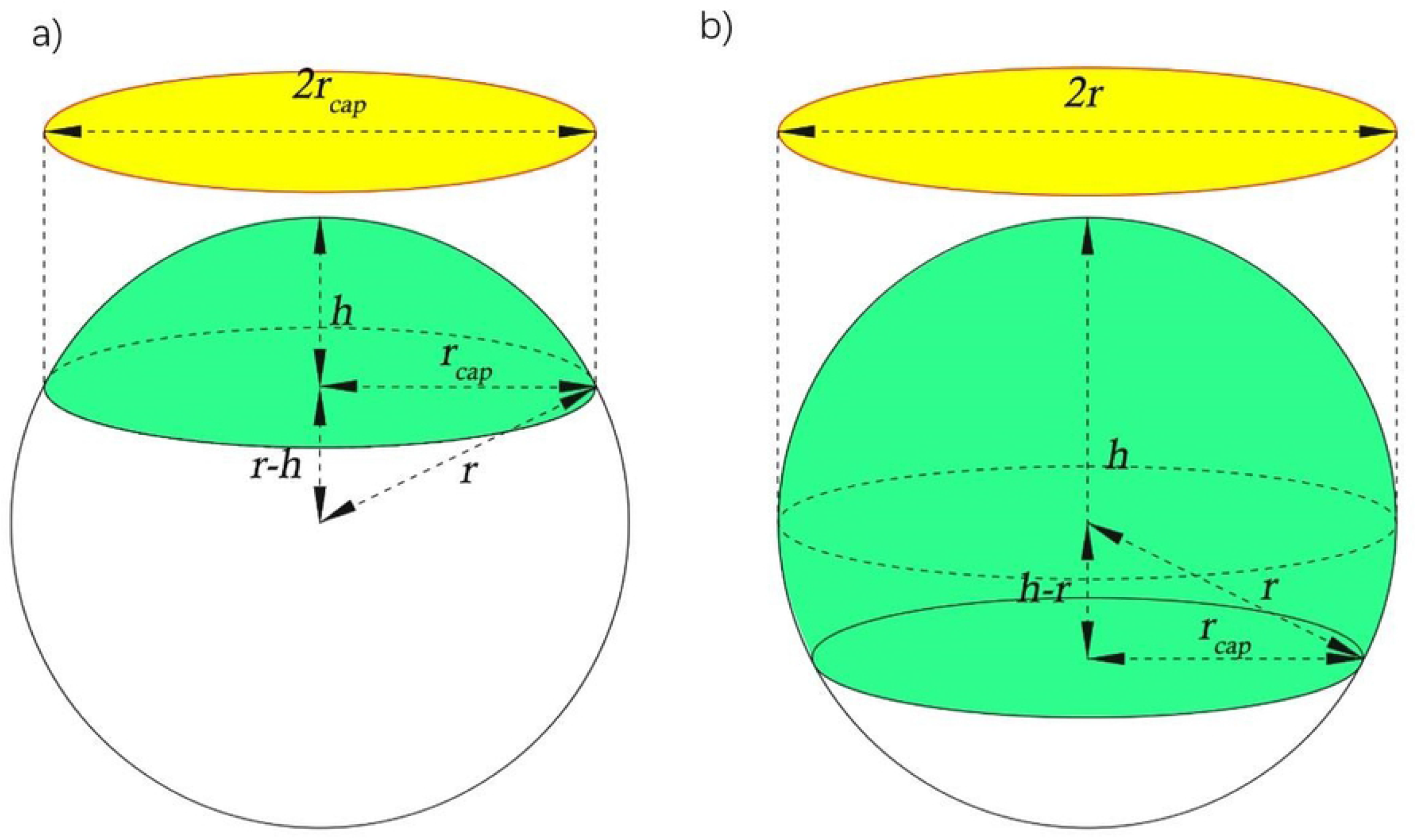
Schematic drawing of top-down imaging drop dimensions for a) hydrophilic and b) hydrophobic surfaces.

Superhydrophobic surfaces are characterised with CA *θ* > 150° [23].

When the radii of the circles in the top-down images are measured, representing the yellow circles in Fig 4, it may be unclear which domain the measurement is in, i.e., whether the maximum visible radius is *r_cap_* or *r*. In practice, the smartphone measures diameter, *d*, and divides by two to get *r_cap_* If the measured drop caplet base radius is *r_cap_*, the volume of the spherical caplet is given by

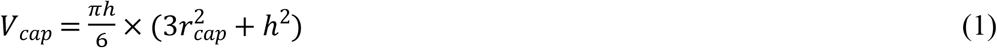

If the volume is known, then using *r_cap_* the height *h* can be calculated. If it is hydrophilic *h* will be less than *r_cap_*. However, if it is larger, then the volume is not correct and instead, the following volume is used to obtained *h* [24]:

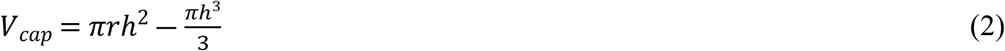

The CA is given in both cases by:

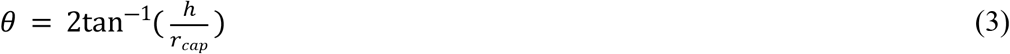

In practice, visual observation can show which regime one is in because the hydrophobic case has a band structure edge rather than a sharp edge, characteristic of two observable diameters, whereas the hydrophilic case only has the caplet base diameter. A flow diagram of the process used is shown in Fig 5.

**Fig 5.**
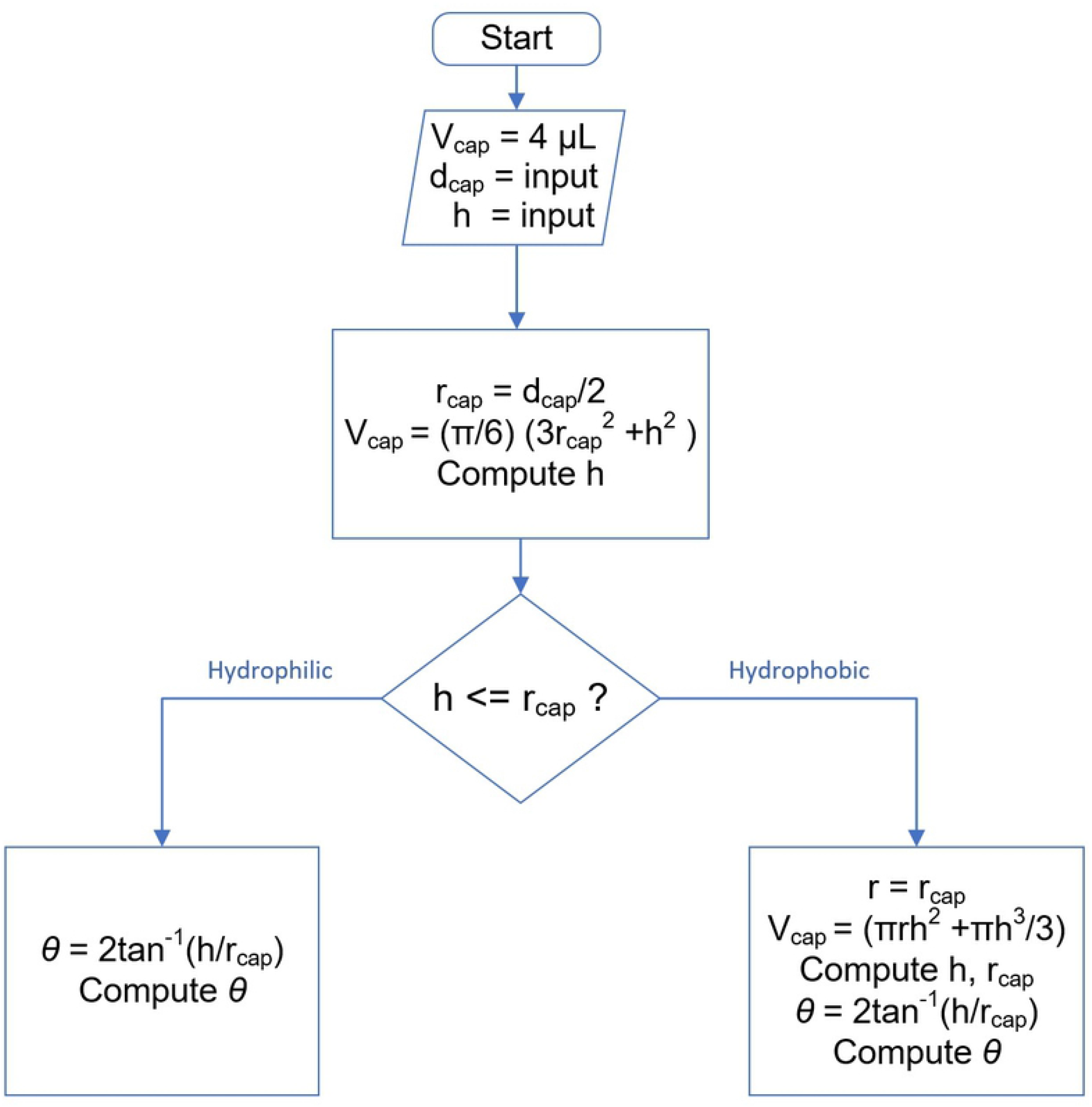
Flow diagram showing the decision-making process to distinguish between hydrophilic and hydrophobic cases.

The advantage of top-down CA measurement is the ability to directly image multiple drops over a surface and therefore map wettability anywhere over that surface. Having this in the field means near real-time measurements are feasible.

### Experimental Validation

To validate the measurement, a piece of tempered glass, similar to Corning Gorilla glass [25], was tested. It is an alkali aluminosilicate sheet glass with a highly compressive thin layer at the surface formed by ion exchange. The glass is said to be hydrophobic because of the fabrication process, which makes it easier to clean. The sample dimensions are *x,y,z* = (70.4 × 153.4 × 0.6) mm. Both top-down and side view results are presented in Fig 6. Side view CA are measured from cut samples and used to benchmark the top-down approach. To reduce or eliminate errors that may be created by gravitational deformation, small water drops of volume *V* = 4 *μ*L are used to standardise these experiments. On the tempered glass, this leads to hydrophobic caplets with spherical diameter *ϕ* = 3.1 ± 0.3 mm.

**Fig 6.**
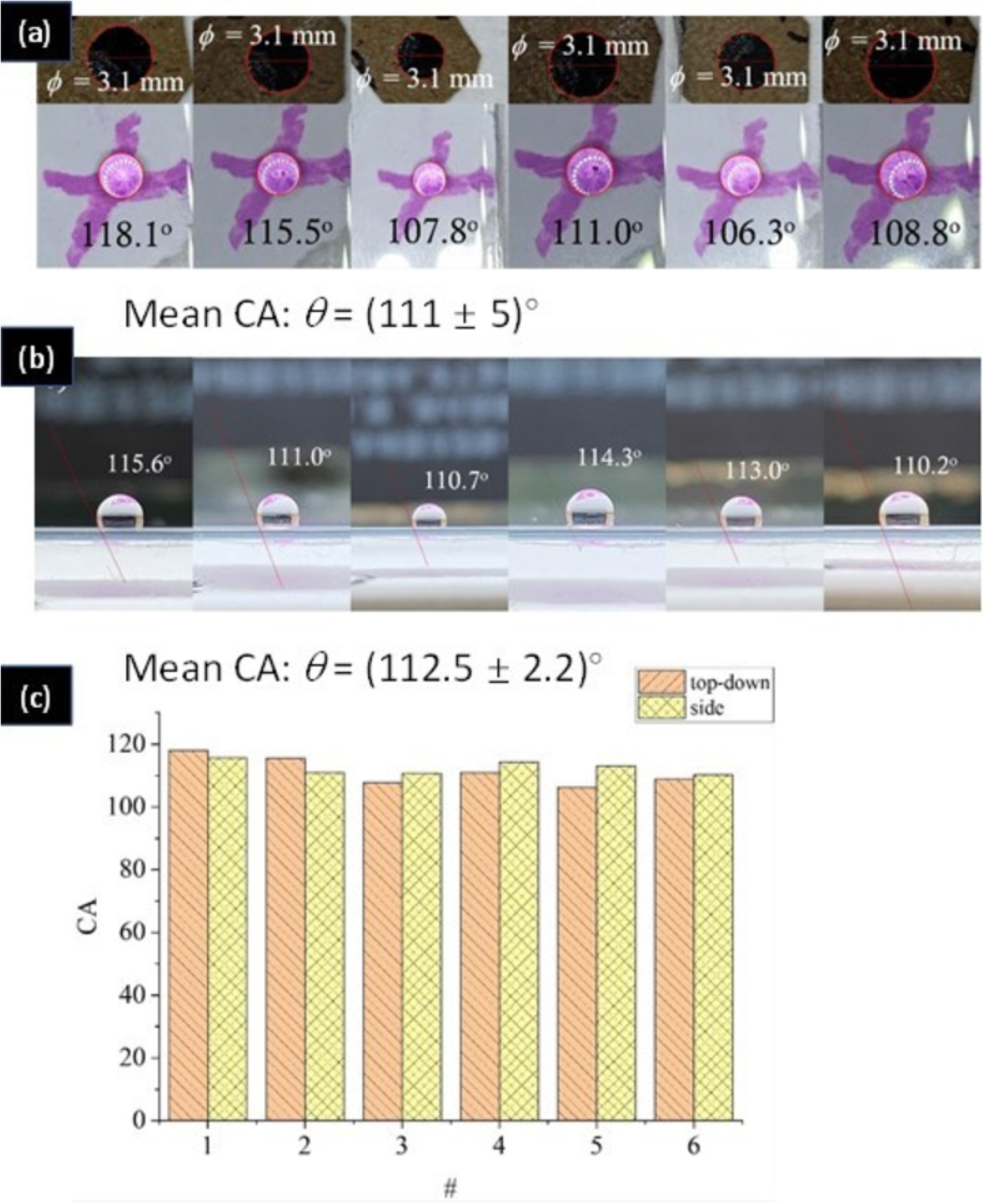
a) Contact angle, CA, measured by the top-down analysis; b) CA measured by side-view analysis using a tangential fit. The mean and standard deviation is shown below the figures a) and b); c) plot of CA versus # corresponding to the sequence of the measured drops in a) and b).

Fig 6 shows that the top-down and side view CA measurement methods produce similar results with low standard deviation demonstrating reliable top-down CA measurements using the smartphone.

### Plant Measurements

To assess the instrument and initially reduce uncertainty from site measurements, sample leaves of Aucuba japonica are gathered from different locations in one place and measured. This ensures identical temperature and environment conditions, helping to identify and assess variations arising from the instrument itself. Times from collection to measurement are *t* = 10 mins. To preserve the initial environment and surface properties of these leaves when fresh and dismiss any effects that can distort surface properties, as well as ensure rapid measurements, the transported samples are kept in sealed bags and the measurements carried out immediately after collection and transport.

Three characteristics are measured and studied with CA mapping.

#### (1) Comparing young and old leaves

Depending on location and planting times, the age of the plants varies and needs to be considered. Young leaves grow at the top of Aucuba japonica, displacing older leaves – these youngest leaves tend to be highest whilst oldest leaves tend to be lowest. Fig 7 summarises the measurements obtained. In general, young leaves are observed to be more hydrophobic than mature ones. This is explained by noting the mature ones are more heavily exposed to environmental variables over a longer time period, including solar radiation, chemicals, wind, temperature changes and rainfall [26,27]. The pH of water can also vary depending on location.

**Fig 7.**
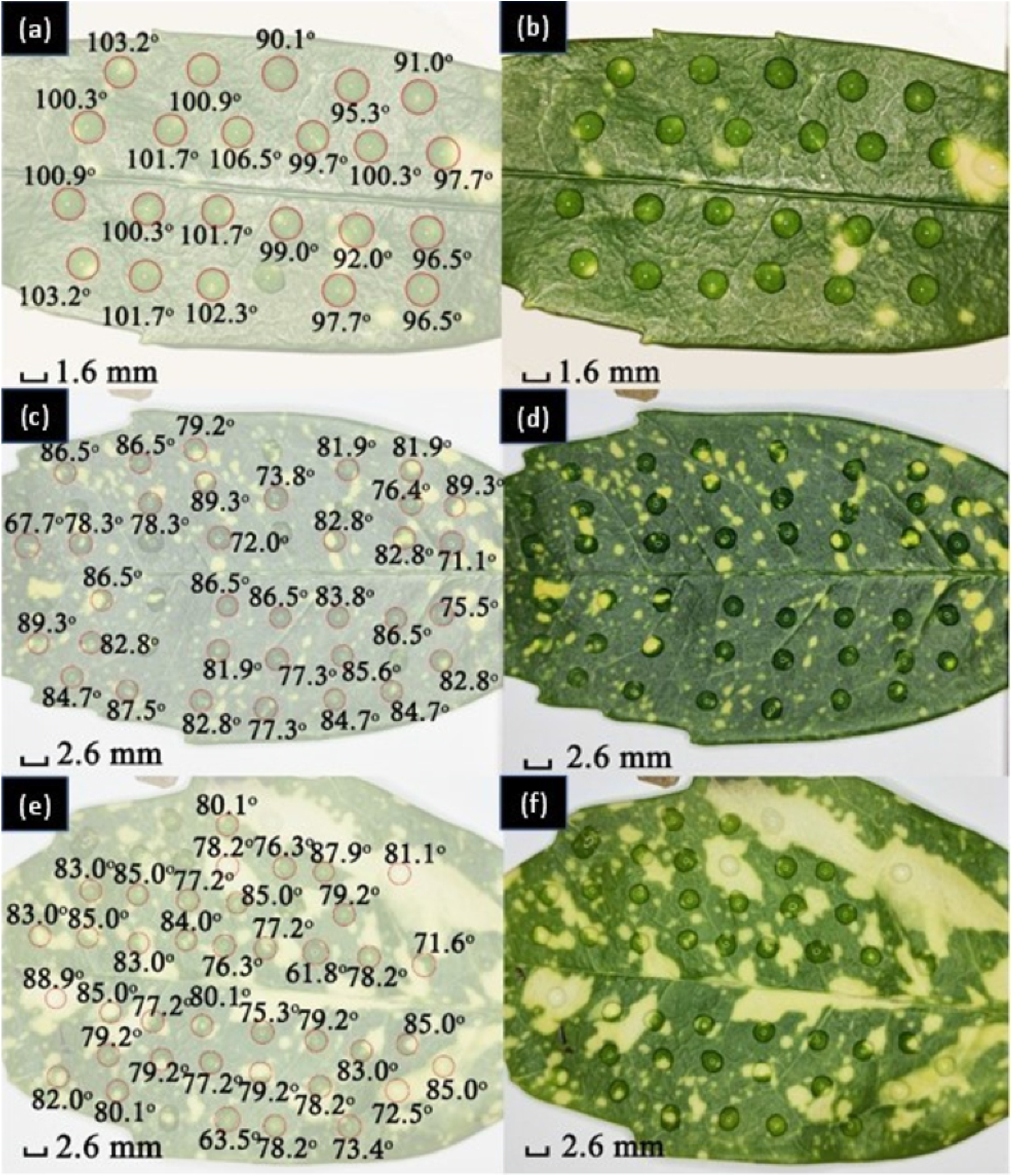
CA measured over the surface of a, b) a young leaf; c, d) a mature leaf; and e, f) a leaf overexposed under sunlight. Images on the left show the measured data overlain on the image of the leaf with drops shown on the right.

The mean CA for the young leaf is *θ* = (99 ± 4)°, and for the mature leaf *θ* = (82 ± 6)°, a notable difference of Δ*θ* ~ 17°. There is an observable change in surface wettability from hydrophobic for a young leaf, to hydrophilic for a mature leaf. The origin of this difference can be attributed to aging over time with exposure to solar radiation and wind in the field.

It was observed that for leaves directly exposed to sunlight compared to those sitting more in shadow without direct solar exposure (Fig 7 e, f)) have a mean CA of *θ* = (79 ± 6)°, which is lower than that of the mature, shadowed plant. That difference can be explained by higher UV exposure and is consistent with the suggestion that it may arise from a leaf that is overexposed to solar radiation for an extended period such that the epicuticular cuticle might be ruined, burnt or partially destroyed [27]. Accelerated aging should, in principle, be induced with exposure to an artificial UV light source something that can be tested. Correlating such a property would provide a novel approach to reliable solar monitoring in the environment over time.

#### (2) Exposure to near UVA light (*λ* = 365 nm)

To explore if solar radiation can accelerate plant aging, or affect the surface properties of Aucuba japonica’s leaves, the leaves were exposed to near UV (UVA) light with wavelength *λ* = 365 nm from a UV diode (power *P* = 6 W, irradiance *I* = (6.75 ~ 7.5) × 10^6^ mW/cm^2^, distance *d* = 9 mm). Fig 8a shows the mean CA for the leaf without UV is *θ* = (91.6 ± 2.8)° and after exposure to UV light for time *t* = 20 min (after 10 minutes, the orientation of leaf would change 180°, total irradiance *I* ~ 1.4 J). The mean CA dropped to *θ* = (90.4 ± 3.0)° (Fig 8b). Fig 8c shows the same leaf exposed to UV radiation for another *t* = 30 min - after 15 min, the orientation of the leaf is rotated by 180° to help ensure even exposure. The total irradiance is *I* ~ 2.2 J. The mean CA changed to *θ* = (86.8 ± 2.5)°, and the leaf on average is hydrophilic, although the standard deviation shows substantial variance. After a much longer continued exposure of *t* = 10 hrs (during the night), the exposure site turned black, indicating that the epidermal microstructures are altered under continued UV radiation. These results show that the leaves of Aucuba japonica will become gradually more hydrophilic over time with solar radiation. This aging process suggests the cuticle layer, made of a hydroxy fatty acid common to many plants, may be slowly ablated until at some point UV light can directly strike the leaf underneath, which then burns. In Aucuba Japonica, the cuticle layer is estimated to be *d* ~ (2 – 3) μm thick, slightly thinner than the cell layer [28]. It is the epidermal layer that is thickest *d* ~ 40 μm. From this reference, the lower cuticle layer at *d* ~ 1 μm is indeed less than the top layer, explaining the observed differences in exposure times.

**Fig 8.**
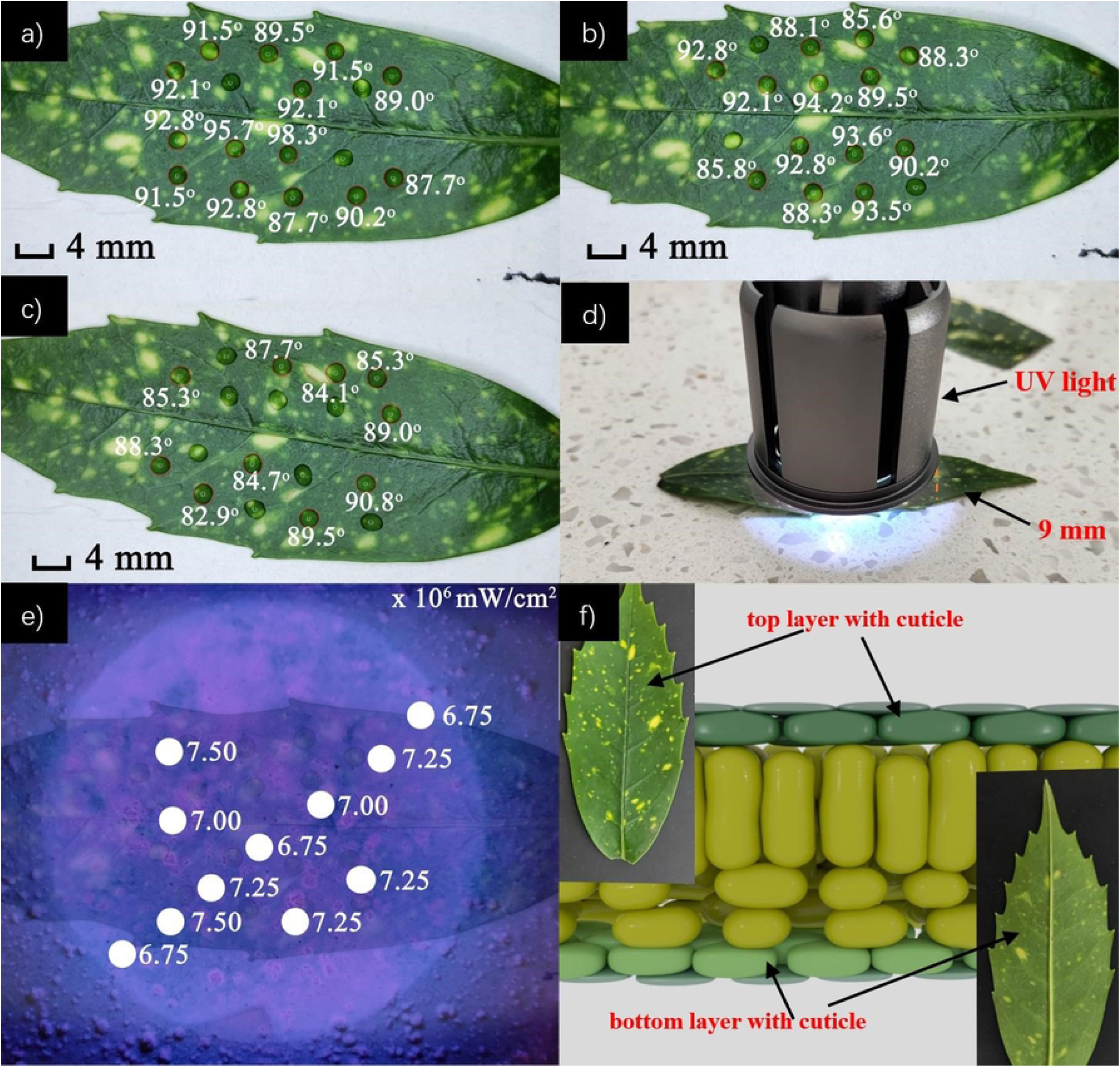
CA and mapping for leaves under UV light: a) CA of the leaf without UVA exposure; b) CA of the leaf after exposure to UVA for *t* = 20 min; c) CA of the leaf after further exposure to UVA for *t* = 30 min; d) shows the UVA (*λ* = 365 nm) source and distance from the sample; and e) shows the area of the leaf under UVA light and the measured irradiance, *I* (mW/cm^2^) at selected points; e) shows the difference between the top layer and bottom layer; and (f) schematic of a typical leaf layer showing typical cuticle layers above and below a leaf [29].

#### (3) The effect of pollution on leaf wettability

To test the potential of this natural sensor for environmental monitoring the impact of pollution on surface wettability is examined. Pollution can drastically affect an environment, and so it would be expected to have an impact on both plant and human health. Consequently, the plant can also act as a monitoring instrument for assessing such pollution. For example, in densely populated cities, particulate matter from cars is a known health problem [30,31]. The concentration of particulate matter will be highest near heavily used traffic. Whether this matter was detectable through CA changes on the leaves of plants growing at and near a street is examined. Figure 2 shows the location chosen.

Compared to previous measurements of leaves away from traffic, three tested leaves (Fig 9 a,b,c) are found to be much more hydrophilic. Mean numbers are *θ* = (58 ± 11)°, *θ* = (52.6 ± 2.5)° and *θ* = (46.7 ± 4.0)°. There is a significant change in CA of Δ*θ* ~ (30 - 40)°, which is consistent with a layer of particulate matter that absorbs and spreads water on the leaf and tries to keep it away [32,33]. This matter stems from the exhaust of vehicles from passing traffic (Fig 2 (b)). Among these leaves, there were poor drop shapes because of ridges on the leaf and other factors where particulates may aggregate in larger quantities. In this case, these drops were not measured as they no longer approximated a spherical caplet and will not be accurately described by eq.’s (1) – (3). Moving away from the street, the surface properties were observed to change with increasing CA of *θ* ~ (88.5 ± 4.7)°, (92.6 ± 2.2)°and (96.0 ± 2.2)°, going back to hydrophobic (Fig 9 d,e,f). Similar results are obtained by lightly brushing the leaves near the road to remove the particulate matter. This dramatic change on CA shows how effective a layer of dust is in protecting the leaf from UV degradation as well as blocking photosynthesis, reducing plant growth.

**Fig 9.**
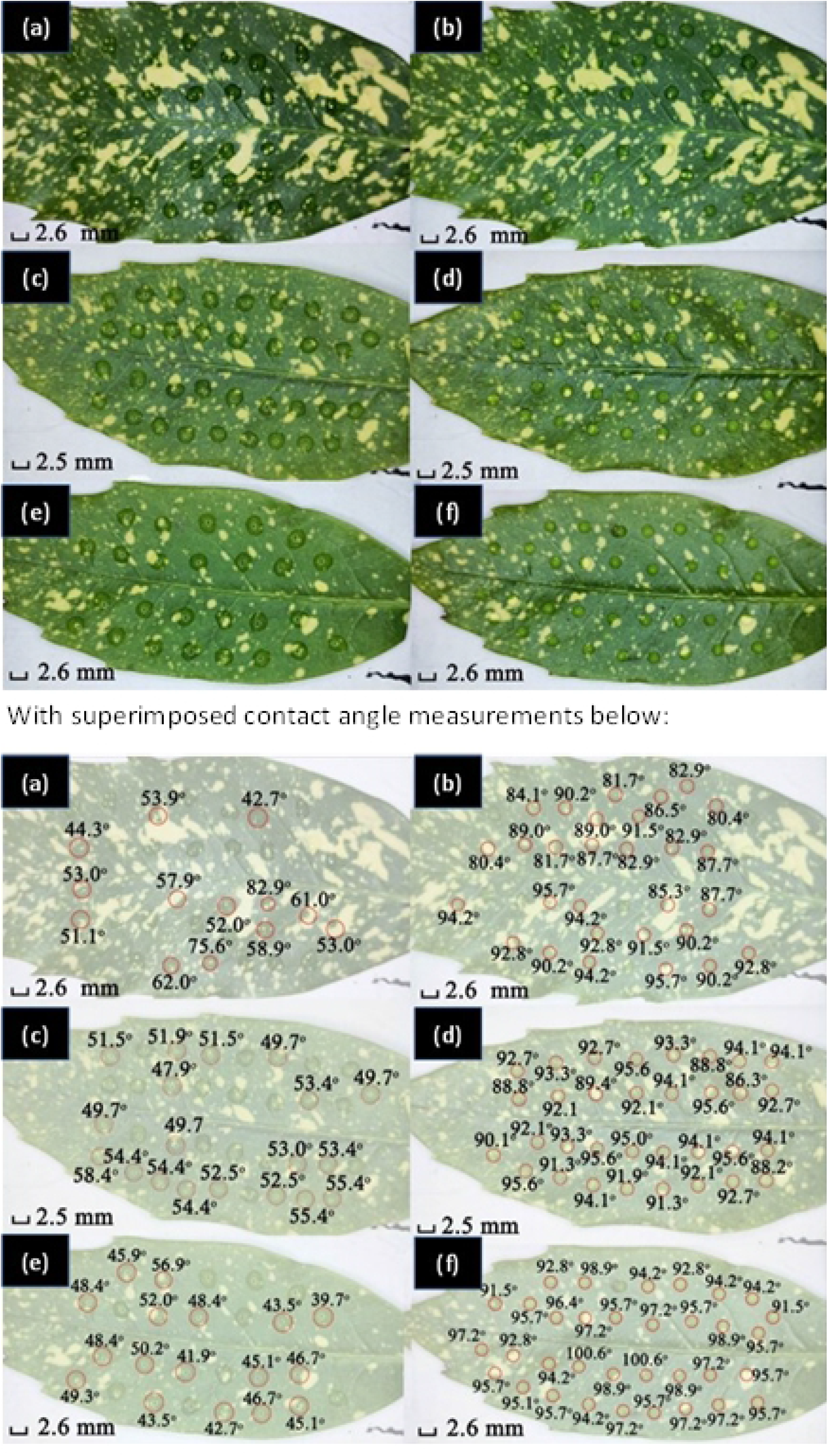
CA mapping for leaves beside the road with heavy traffic (refer to Figure 1). Various leaf images are shown. Top: a), c), e) show calibrated drops on leaves without brushing, whilst b), d), f) show calibrated drops on leaves after brushing. Further, a), b) are for a leaf under sunlight without shelter, whereas c), d), e), f) are for leaves near the concrete pillar in Figure 1. All the bottom images below show the location and measured contact angles overlain on the top images.

## Discussion and Conclusion

Smartphone-based top-down CA mapping measurements are effective as a tool that can be taken close to the sensor site in the field and used to collect data, making near real time measurements feasible. The quality of the measurements is competitive with side-based measurements that rely on more expensive laboratory bench equipment but have the advantage of ease of measurement and portability. This offers a convenient, reliable way to measure surface wettability on-site for complex samples such as leaves. In addition, the top-down approach allows an entire surface to be easily mapped - this translation of a smart IoT tool into the field enables reliable measurement and mapping of properties of leaves in near real-time, illustrating the value of plants as intrinsic environmental sensors. For botanists, this technology presents a scientific breakthrough enabling large data generation that can provide reliable information about plant evolution and the environment it is in.

Combining smartphones with natural plant sensors is a hybrid technology that opens a novel approach to environmental monitoring, the basis of a bionic internet-of-things (*b*-IoT). Here, specifically, the use of Aucuba japonica, a robust plant that has wide regional distribution, has been characterised and significant changes in parameters were detected that make such bionic sensor networks feasible. Very highly sensitive detection of particulate matter deposited by the environment on the leaf was possible, offering large dynamic sensor range. The long-term performance of the sensors compares favorably with the need for continual maintenance required with electronic versions. To improve accuracy and reliability over longer time periods, changes arising from aging and impact need to be understood and calibrated into analysis. For example, young leaves are a little hydrophobic while mature ones are slightly hydrophilic, reflecting a degree of aging in the environment attributed to the degradation of the leaf cuticle over time. The role of UV exposure in aging the sensors was explored and confirmed, providing a novel tool for recording and tracking solar measurements over time. Degradation of the protective waxy layer occurs before the plant itself is exposed and destroyed. It was found that under the leaves, which do not require access to the sun and are less exposed, the cuticle layer is likely to be much thinner and therefore, removal is faster where the epidermal and cell layers burn earlier. The conjecture on different thicknesses at the top and bottom of the leaves is supported by existing literature that has measured this [28], confirming the validity of the measurements here. In fact, the difference in layers is partly responsible for the way leaves turn outwards to collect sunlight, effectively a multilayer thin film stress effect.

This form of bionic instrumentation is particularly attractive because it couples low cost, low maintenance, aesthetically pleasing and naturally growing sensors with smartphone diagnostics. In principle, other technologies could be integrated into the *b*-IoT, including for example, bionic jellyfish or scorpion skins which mimic natural organisms [34,35]. Here, we directly integrate living organisms though in principle the *b*-IoT can use both. They blend into an environment, creating aesthetic, health and wellness improvements for people and animals, a feature that contrasts strongly with current IoT sensor technologies and networks. Aucuba Japonica is an example that is regionally widespread offering a sensing tool that can be used to map responses across the globe, all collated at one point. Through the internet, the sensor data from each plant can be brought together so that mapping across regions where potentially thousands of routine measurements are taken in near real-time. They are linked through the same cloud diagnostics into which each edge computer communicates. A smart tool such as a contact angle analyser on a phone means that this technology is widely accessible to the consumer, perfect for citizen engagement in a networked super diagnostic for environmental monitoring and, by extension, potentially for climate change analysis. The totality of the integrated plant-internet system marks a first step towards a *b*-IoT, a crucial technology development that can directly monitor environmental impact through several causes, including natural and human-induced impact. It can potentially expand the scope of tools in the assessment of climate change more broadly whilst simultaneously enhancing human health and wellbeing.

## Acknowledgment

Y. Guo acknowledges an Australian International Research Training Program Scholarship (IRTP, Department of Education and Training, Australia) under Prof Canning. Private funding is also acknowledged.

## Reference

[1] Hossain MA, Brito-Rodriguez B, Sedger LM, Canning J. A cross-disciplinary view of testing and bioinformatic analysis of SARS-CoV-2 and other human respiratory viruses in pandemic settings. IEEE Access 2021:1–1. https://doi.org/10.1109/ACCESS.2021.3133417.

[2] Cammers-Goodwin SI. Revisiting Smartness in the Smart City. The Oxford Handbook of Philosophy of Technology 2022;9:169.

[3] Hall C, Knuth M. An Update of the Literature Supporting the Well-Being Bene¢ts of Plants: A Review of the Emotional and Mental Health Bene¢ts of Plants 2019:9.

[4] Ohi T, Kajita T, Murata J. Distinct geographic structure as evidenced by chloroplast DNA haplotypes and ploidy level in Japanese *Aucuba* (Aucubaceae). Am J Bot 2003;90:1645–52. https://doi.org/10.3732/ajb.90.11.1645.

[5] Zeng X, Guo F, Ouyang D. A review of the pharmacology and toxicology of aucubin. Fitoterapia 2020;140:104443. https://doi.org/10.1016/j.fitote.2019.104443.

[6] Bean. Trees and shrubs hardy in the British Isles, edn 8, I. A-C. London: John Murray; 1970.

[7] Tachibana S, Watkins C. Botanical Transculturation: Japanese and British Knowledge and Understanding of Aucuba japonica and Larix leptolepis 1700–1920. Environment and History n.d.:30.

[8] Spotted Laurel · Plant Finder n.d. https://www.smartwatermark.org/smartwateradvice/plant-finder/plant/spotted-laurel/ (accessed December 23, 2021).

[9] Uke A, Pinili MS, Natsuaki KT, Geering ADW. Complete genome sequence of aucuba ringspot virus. Arch Virol 2021;166:1227–30. https://doi.org/10.1007/s00705-021-04977-4.

[10] Korsakova S, Plugatar Yu, Ilnitsky O, Karpukhin Yu. A research on models of the photosynthetic light response curves on the example of evergreen types of plants 2019:1.163Mb. https://doi.org/10.15159/AR.19.065.

[11] Ginetti B, Ragazzi A, Carmignani S, Moricca S. Collar Rot and Crown Wilting by Phytophthora pachypleura on Aucuba japonica in Italian Nurseries. Plant Disease 2015;99:1860–1860. https://doi.org/10.1094/PDIS-01-15-0120-PDN.

[12] Lomtatidze N, Alasania N, Machutadze E. Bioecological peculiarities of introduced exotic species of Japanese laurel (Aucuba japonica Thunb) at the Black Sea coast of Ajara, Georgia. Bulletin of the Georgian National Academy of Sciences 2013;7:110–3.

[13] Aucuba japonica. Wikipedia 2021.

[14] Search n.d. https://www.gbif.org/species/3033077 (accessed December 29, 2021).

[15] Liu Z, Hua Q, Wang J, Liang Z, Zhou Z, Shen X, et al. Prussian blue immunochromatography with portable smartphone-based detection device for zearalenone in cereals. Food Chemistry 2022;369:131008. https://doi.org/10.1016/j.foodchem.2021.131008.

[16] Biswas PC, Rani S, Hossain MA, Islam MR, Canning J. Recent Developments in Smartphone Spectrometer Sample Analysis. IEEE J Select Topics Quantum Electron 2021;27:1–12. https://doi.org/10.1109/JSTQE.2021.3075074.

[17] Drelich JW. Contact angles: From past mistakes to new developments through liquid-solid adhesion measurements. Advances in Colloid and Interface Science 2019;267:1–14. https://doi.org/10.1016/j.cis.2019.02.002.

[18] Wang H, Shi H, Wang Y. The Wetting of Leaf Surfaces and Its Ecological Significances. In: Aliofkhazraei M, editor. Wetting and Wettability, London: IntechOpen; 2015, p. 295–321. https://doi.org/10.5772/61205.

[19] Neufeld HS, Jernstedt JA, Haines BL. Direct foliar effects of simulated acid rain: I. Damage, growth and gas exchange. New Phytol 1985;99:389–405. https://doi.org/10.1111/j.1469-8137.1985.tb03667.x.

[20] Burkhardt J. Hygroscopic particles on leaves: nutrients or desiccants? Ecological Monographs 2010;80:31.

[21] Kardel F, Wuyts K, Babanezhad M, Wuytack T, Adriaenssens S, Samson R. Tree leaf wettability as passive bio-indicator of urban habitat quality. Environmental and Experimental Botany 2012;75:277–85. https://doi.org/10.1016/j.envexpbot.2011.07.011.

[22] Bigelow WC, Pickett DL, Zisman WA. Oleophobic monolayers: I. Films adsorbed from solution in non-polar liquids. J Colloid Sci 1946;1:513–38. https://doi.org/10.1016/0095-8522(46)90059-1.

[23] Dutra G, Canning J, Padden W, Martelli C, Dligatch S. Large area optical mapping of surface contact angle. Opt Express 2017;25:21127. https://doi.org/10.1364/OE.25.021127.

[24] Janeczko C, Martelli C, Canning J, Dutra G. Assessment of Orchid Surfaces Using Top-Down Contact Angle Mapping. IEEE Access 2019;7:31364–75. https://doi.org/10.1109/ACCESS.2019.2902730.

[25] CORNING® GORILLA® GLASS | THE TECHNOLOGY 2013. https://web.archive.org/web/20131105090231/http://www.corninggorillaglass.com/technology (accessed January 10, 2022).

[26] Rykaczewski K, Jordan JS, Linder R, Woods ET, Sun X, Kemme N, et al. Microscale Mechanism of Age Dependent Wetting Properties of Prickly Pear Cacti (*Opuntia*). Langmuir 2016;32:9335–41. https://doi.org/10.1021/acs.langmuir.6b02173.

[27] Fernández V, Sancho-Knapik D, Guzmán P, Peguero-Pina JJ, Gil L, Karabourniotis G, et al. Wettability, Polarity, and Water Absorption of Holm Oak Leaves: Effect of Leaf Side and Age. Plant Physiol 2014;166:168–80. https://doi.org/10.1104/pp.114.242040.

[28] Onoda Y, Schieving F, Anten NPR. A novel method of measuring leaf epidermis and mesophyll stiffness shows the ubiquitous nature of the sandwich structure of leaf laminas in broad-leaved angiosperm species. Journal of Experimental Botany 2015;66:2487–99. https://doi.org/10.1093/jxb/erv024.

[29] Zhang Q, Zhang M, Ding Y, Zhou P, Fang Y. Composition of photosynthetic pigments and photosynthetic characteristics in green and yellow sectors of the variegated Aucuba japonica ‘Variegata’ leaves. Flora 2018;240:25–33. https://doi.org/10.1016/j.flora.2017.12.010.

[30] Yang Q, Shen H, Liang Z. Analysis of particulate matter and carbon monoxide emission rates from vehicles in a Shanghai tunnel. Sustainable Cities and Society 2020;56:102104. https://doi.org/10.1016/j.scs.2020.102104.

[31] Giechaskiel B, Maricq M, Ntziachristos L, Dardiotis C, Wang X, Axmann H, et al. Review of motor vehicle particulate emissions sampling and measurement: From smoke and filter mass to particle number. Journal of Aerosol Science 2014;67:48–86. https://doi.org/10.1016/j.jaerosci.2013.09.003.

[32] Jagels R. Leaf Wettability as a Measure of Air Pollution Effects. In: Percy KE, Cape JN, Jagels R, Simpson CJ, editors. Air Pollutants and the Leaf Cuticle, Berlin, Heidelberg: Springer Berlin Heidelberg; 1994, p. 97–105. https://doi.org/10.1007/978-3-642-79081-2_7.

[33] Cape JN, Paterson IS, Wolfenden J. Regional variation in surface properties of Norway spruce and scots pine needles in relation to forest decline. Environmental Pollution 1989;58:325–42. https://doi.org/10.1016/0269-7491(89)90143-7.

[34] Vogel V. Bionic jellyfish. Nature Mater 2012;11:841–2. https://doi.org/10.1038/nmat3438.

[35] Zhiwu H, Junqiu Z, Chao G, Li W, Ren L. Erosion Resistance of Bionic Functional Surfaces Inspired from Desert Scorpions. Langmuir 2012;28:2914–21. https://doi.org/10.1021/la203942r.

